# Improving the classification of neuropsychiatric conditions using gene ontology terms as features

**DOI:** 10.1101/393082

**Authors:** Thomas P. Quinn, Samuel C. Lee, Svetha Venkatesh, Thin Nguyen

## Abstract

Although neuropsychiatric disorders have a well-established genetic background, their specific molecular foundations remain elusive. This has prompted many investigators to design studies that identify explanatory biomarkers, and then use these biomarkers to predict clinical outcomes. One approach involves using machine learning algorithms to classify patients based on blood mRNA expression from high-throughput transcriptomic assays. However, these endeavours typically fail to achieve the high level of performance, stability, and generalizability required for clinical translation. Moreover, these classifiers can lack interpretability because informative genes do not necessarily have relevance to researchers. For this study, we hypothesized that annotation-based classifiers can improve classification performance, stability, generalizability, and interpretability. To this end, we evaluated the performance of four classification algorithms on six neuropsychiatric data sets using four annotation databases. Our results suggest that the Gene Ontology Biological Process database can transform gene expression into an annotation-based feature space that improves the performance and stability of blood-based classifiers for neuropsychiatric conditions. We also show how annotation features can improve the interpretability of classifiers: since annotation databases are often used to assign biological importance to genes, annotation-based classifiers are easy to interpret because the biological importance of the features are the features themselves. We found that using annotations as features improves the performance and stability of classifiers. We also noted that the top ranked annotations tend contain the top ranked genes, suggesting that the most predictive annotations are a superset of the most predictive genes. Based on this, and the fact that annotations are used routinely to assign biological importance to genetic data, we recommend transforming gene-level expression into annotation-level expression prior to the classification of neuropsychiatric conditions.

## 1 Introduction

The field of neuropsychiatry involves a collection of complex heterogeneous disorders with a known social and genetic aetiological basis. However, the specific molecular and genetic foundations of these disorders remain elusive. This has prompted a number of genomic studies which seek to identify the transcriptomic and genetic signatures associated with each disorder. As a corollary to this work, investigators have sought to use genomic data to build classifiers that can accurately predict neuropsychiatric conditions. To this end, some studies have used mRNA expression, measured in the blood as biomarkers, to predict neuropsychiatric disorders like autism [10, 15, 35], as previously done for cancer [11, 1]. Since blood is easy to collect, an accurate blood-based classifier could have direct clinical utility. Although neuropsychiatric disorders are traditionally described as disorders of the brain, they are hypothesised to involve systemic processes [22], and blood has been shown to serve as a useful approximation for what happens in the brain [32].

The classification of complex heterogeneous disorders using transcriptomic data is difficult. Such data tend to have properties that pose a challenge to building accurate and generalizable classifiers. First, these data are high-dimensional with many more features than samples (the “*p >> N* problem”). Second, individual features tend to have small explanatory effect sizes. Third, the binary class labels used to differentiate “cases” from “controls” often describe a vastly heterogeneous phenotype. Fourth, there exists between-study differences in cohort demographics that compound the limitations already imposed by small sample sizes. Most often, the problem of high-dimensionality is addressed using *feature selection* (whereby features are prioritized by external or embedded methods). However, it is also possible to reduce feature space through *feature engineering* (a process which uses domain-specific knowledge to transform the measured feature space into a new feature space). Databases that map gene symbols to functional annotations typically have fewer annotations than annotated genes, and therefore offer a systematic way to perform feature engineering (as used successfully in the classification of cancer [8, 39]).

The use of annotations for feature engineering is not without challenges. For example, there are many annotation databases available, all of which provide abstractions of biology that might not capture the full complexity of biological systems. Even if they do, the curation of these databases are ongoing endeavours, and therefore not necessarily exhaustive. Moreover, it remains an open question as to how best to engineer a gene-level feature space into an annotation-level feature space. Yet, despite these challenges, we hypothesize that annotation-based feature engineering would add value to classification. First, it ameliorates the “*p >> N* problem” (thus potentially improving classifier performance). Second, it aggregates individual signals into a higher-order abstraction such that small effect sizes may accumulate and cohort differences may converge (thus potentially improving classifier stability and generalizability). Finally, these annotations, as abstractions of biology, reflect domain-specific knowledge with a clear meaning to biological scientists (thus improving classifier interpretability).

There are two general approaches to developing an expert-curated annotation database. The first associates gene with biochemical pathways based on experimental evidence. The second associates genes with phenotypes or disease states. However, with several databases available, and a number of ways to engineer features based on these databases, it is useful to assess empirically which approaches to annotation-based feature engineering, if any, work best. In this study, we assess whether annotation-based feature engineering maintains or improves performance across 6 neuropsychiatric data sets, as benchmarked using 4 classification algorithms and 4 annotation databases. In addition, we measure whether annotation-based feature engineering improves the stability and generalizability of the classifiers. Our results show that annotation-based classifiers outperform gene-based classifiers in terms of accuracy and stability, and that the most predictive annotations contain the most predictive genes. We conclude by discussing how the use of annotations can improve the interpretability of a classifier because they represent the data in a space that captures how scientists conceptualize biology.

## 2 Methods

### 2.1 Data acquisition

We acquired six blood-based microarray data sets from the NCBI Gene Expression Omnibus (GEO), all relevant to the field of neuropsychiatry. Three data sets compare the whole blood (GSE18123-GPL570; GSE18123-GPL6244 [15]) and lymphocytes (GSE25507 [2]) of autism spectrum disorder (ASD) patients with typically developing (TD) controls. Another, GSE38484, compares the whole blood of schizophrenic patients with controls [7]. The fifth, GSE98793, compares the whole blood of major mood disorder (MDD) patients with controls [18]. The sixth, GDS5393, compares the peripheral blood of bipolar I patients before and after lithium treatment [5]. Data were acquired already normalized and were not modified further.

### 2.2 Data preparation

Source microarray data measure transcript expression using probes. We converted probe-level expression to gene-level measurements by aggregating mapped values by a median summary. For probe-level to gene-level mapping, we used the appropriate AnnotationDbi package [23].

### 2.3 Feature engineering

We then converted gene-level measurements to annotation-level measurements by aggregating mapped values using one of four methods: mean, median, sum, and variance. For this, we used the Gene Ontology Biological Process (**BP**) and Molecular Function (**MF**) [3, 34], Disease Ontology (**DO**) [13], and Human Phenotype Ontology (**HPO**) [14] databases. We only included annotations that map to at least 5 genes, and did not exclude any probes, genes, or annotations due to mapping ambiguities. For the three ASD data sets, we only included genes (and annotations) represented across all microarray platforms.

### 2.4 Classification

We performed all machine learning using the R package exprso, a software tool that wraps complete machine learning pipelines for use in a high-throughput manner [26]. For this analysis, we trained binary classification models, defining class labels as “control” versus other.

We applied two machine learning pipelines. The first measures within-study performance. The second measures across-study performance. These pipelines differ only in how the training set is defined and how classifier performance is calculated. Note that, in all cases, we selected features and built models on a training set that is separate from the test set, making the test set statistically independent.

#### 2.4.1 Feature input

For each of the six microarray data sets, we represented transcript expression at the gene-level or at one of four annotation-levels. These annotation-level measurements are created from the gene-level measurements using one of four summary methods (described above). Each unique feature space contributes a unique data set upon which we applied the machine learning pipelines.

#### 2.4.2 Training set split

For the within-study pipeline, all training sets contain a stratified random sample of the data, balanced by class label. This approach ensures that both the training and test sets have an equal number of cases and controls. Each training set has 67% of the balanced data. For the across-study pipeline, the training and test sets are separate microarray data sets.

#### 2.4.3 Feature selection

For all pipelines, we selected gene-level and annotation-level features from the training set using Student’s *t*-test [30]. We also selected features by random sampling to provide a point of reference.

#### 2.4.4 Model building

For all pipelines, we trained a model on the training set using a decision tree (DT) (via rpart::rpart [33]), logistic regression (LR) (via stats::glm), random forest (RF) (via randomForest::randomForest [20]), or support vector machine (SVM) (via e1071::svm [21]), with the top *N* = [2, 3*, ...,* 64, 128] selected features. For all implementations, we use the default arguments except when building decision trees (for which we prune with the argument cp = 0.2). For the purpose of this study, we do not tune hyper-parameters.

#### 2.4.5 Estimating performance

For the within-study pipeline, we calculated classifier performance on a single data set by repeating the training set split, feature selection, and classifier construction procedure *B* = 100 times. This allows us to calculate a stable expected (i.e., average) accuracy. In the literature, this approach is sometimes called “Monte Carlo cross-validation” [24].

For the across-study pipeline, we calculated the performance of a classifier built on one data set and then deployed on another. We did this by treating the entirety of the source data set as the training set and the entirety of the target data set as the test set. As such, we applied feature selection and classifier construction only once per data set pair.

### 2.5 Measuring feature stability

In this paper, we define stability as the likelihood that the same features would get selected from two separate training set splits of the same source data. For each data set and feature space, we measure the stability of feature selection across *B* = 100 training sets using two methods.

The first calculates the average Baroni-Urbani and Buser (BUB) Overlap [4] for each pair of the 100 ranked features (analogous to the Rand Index [27]):

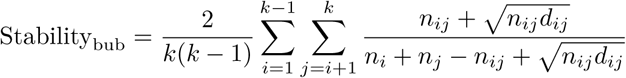

where *k* is the number of training sets, *n*_*i*_ is the number of features selected in training set *i*, *n*_*ij*_ is the number of features selected in training sets *i* and *j*, and *d*_*ij*_ is the number of features not selected in training sets *i* or *j*.

For a given feature space *f* (gene- or annotation-level), Stability_bub_ approaches 1 as the number of selected features (*n*_*f*_) approaches the total number of features (*N*_*f*_). Although Stability_bub_ depends on *N*_*f*_ by definition, it has an equivalent null distribution for a fixed number of features as a percentile of the total number of features 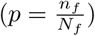 (see Supplemental Figures). Therefore, we can compare gene- and annotation-level feature selection for any percentile *p* of top ranked features to determine whether one feature space is more stable than another, *independent of its size*.

The second calculates the average Spearman’s Rank Correlation Coefficient [29] for each pair of the 100 ranked features:

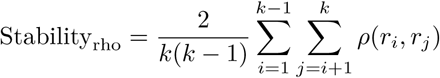

where *k* is the number of training sets, *ρ* is Spearman’s Rank Correlation and *r*_*i*_ is the ranked list of selected features for training set *i*.

Visualizations of stability show all *k*(*k −* 1)*/*2 instances, not the average.

### 2.6 Measuring generalizability

We assess generalizability through the across-study pipeline, wherein the training and test sets are separate microarray data sets. Two of these are independent ASD data sets collected as part of the same larger study (GSE18123). The third is an ASD data set from another study (GSE25507).

### 2.7 Measuring information capture

>Each annotation-level measurement is calculated by aggregating across a set of gene-level measurements. Feature selection reduces the total feature space to a subset of annotations. We define a “gene member set” as the set of the genes which correspond to a subset of annotations. We define “information capture” as the extent to which important annotation-based features contain the important gene-based features in their “gene member set”.

To quantify the “information capture” for an annotation, we compared the “gene member set” for each subset of annotations (sized *N* = [2, 3*, …,* 64, 128]) with its corresponding gene set. Specifically, we used the Fisher Exact Test to test the null hypothesis that a gene set and a “gene member set” are not jointly selected (one-tailed). We performed this test for each ASD data set, feature space, and number of features selected. Since the three ASD data sets contain the same “gene universe”, results are comparable across data sets.

For the Fisher Exact Test, large odds ratios (ORs) suggest that the top selected annotations contain the top selected genes, and that the respective annotation-based classifier would capture the same information as its corresponding gene-based classifier. When comparing ORs with classifier performance, we used the average intra-study validation accuracy across bootstraps (i.e., for each classifier size, considering SVM classifiers only).

## 3 Results

### 3.1 Annotations as features improve performance

We evaluated the performance of four classification algorithms on six neuropsychiatric data sets using five feature spaces. Of these, four feature spaces were annotation-level summaries of gene-level expression. We repeated this by aggregating gene expression based on four summary methods (mean, median, sum, and variance), and compared the resultant cross-validation accuracies. We found that, across all bootstrapped combinations of classification algorithms, classifier sizes, feature spaces, and data sets, mean-based and sum-based summaries performed marginally better than median-based and variance-based summaries (*p < .*05, see Table 1). As such, all subsequent analyses, tables, and figures use mean-based summaries only.

**Table 1:**
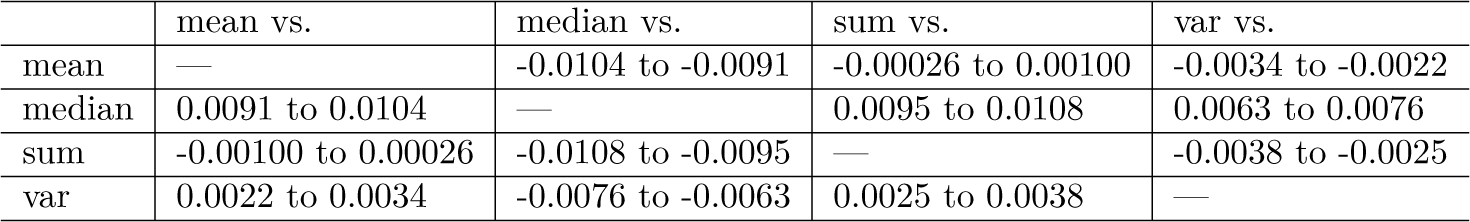
This table shows the 95% confidence intervals for the mean differences of classifier performance between annotation-based classifiers aggregated by mean, median, sum, or variance summary statistics. We found that, across all bootstrapped combinations of classification algorithms, classifier sizes, feature spaces, and data sets, mean-based and sum-based summaries performed marginally better than median-based and variance-based summaries.

|  | mean vs. | median vs. | sum vs. | var vs. |
| --- | --- | --- | --- | --- |
| mean | — | -0.0104 to -0.0091 | -0.00026 to 0.00100 | -0.0034 to -0.0022 |
| median | 0.0091 to 0.0104 | — | 0.0095 to 0.0108 | 0.0063 to 0.0076 |
| sum | -0.00100 to 0.00026 | -0.0108 to -0.0095 | — | -0.0038 to -0.0025 |
| var | 0.0022 to 0.0034 | -0.0076 to -0.0063 | 0.0025 to 0.0038 | — |

Next, we found that, across all bootstrapped combinations of classifier sizes, feature spaces, and data sets, support vector machines (SVM) were the highest performing classification algorithm (*p < .*05 by *t*-test, see Table 2), while decision trees (DT) were the worst. We also found that, across all bootstrapped combinations of classifier sizes and data sets, training SVMs with the **BP** feature space outperformed all other feature spaces (*p < .*05 by *t*-test, see Table 3). This also holds true across for the other classification algorithms (data in Supplement). Yet, in all cases, performance gains are marginal, with each optimal choice adding only about 1% to validation accuracies.

**Table 2:** This table shows the 95% confidence intervals for the mean differences of classifier performance between mean-based classifiers built with a logistic regression (LR), decision tree (DT), random forest (RF), or support vector machine (SVM). We found that, across all bootstrapped combinations of classifier sizes, feature spaces, and data sets, SVMs were the highest performing classification algorithm.

|  | buildLR vs. | buildDT vs. | buildRF vs. | buildSVM vs. |
| --- | --- | --- | --- | --- |
| buildLR | — | -0.069 to -0.067 | 0.025 to 0.027 | 0.027 to 0.030 |
| buildDT | 0.067 to 0.069 | — | 0.093 to 0.095 | 0.096 to 0.098 |
| buildRF | -0.027 to -0.025 | -0.095 to -0.093 | — | 0.0019 to 0.0041 |
| buildSVM | -0.030 to -0.027 | -0.098 to -0.096 | -0.0041 to -0.0019 | — |

**Table 3:** This table shows the 95% confidence intervals for the mean differences of classifier performance between mean-based and SVM-based annotation-level classifiers (built with the Gene Ontology Biological Process (**BP**) and Molecular Function (**MF**), Disease Ontology (**DO**), and Human Phenotype Ontology (**HPO**) databases), and gene-level classifiers. We found that, across all bootstrapped combinations of classifier sizes and data sets, training SVMs with the **BP** feature space outperformed all other feature spaces.

|  | bp vs. | do vs. | gene vs. | hpo vs. | mf vs. |
| --- | --- | --- | --- | --- | --- |
| bp | — | -0.016 to -0.011 | -0.016 to -0.011 | -0.027 to -0.022 | -0.0117 to -0.0068 |
| do | 0.011 to 0.016 | — | -0.0025 to 0.0023 | -0.014 to -0.009 | 0.0015 to 0.0065 |
| gene | 0.011 to 0.016 | -0.0023 to 0.0025 | — | -0.0140 to -0.0089 | 0.0017 to 0.0066 |
| hpo | 0.022 to 0.027 | 0.009 to 0.014 | 0.0089 to 0.0140 | — | 0.013 to 0.018 |
| mf | 0.0068 to 0.0117 | -0.0065 to -0.0015 | -0.0066 to -0.0017 | -0.018 to -0.013 | — |

Figure 1 shows the average performance of SVM classifiers, per classifier size, for each combination of data set and feature space (features selected by *t*-test). Visually, we see that the **BP** tends to perform among the best, except possibly for one ASD data set (GSE18123-GPL570). Figure 2 projects these same data as a box plot (with each point representing a different classifier size). These data also show that the **BP** space tends to perform as well as (or better than) others, although there is variability across the six neuropsychiatric data sets. We refer the reader to the Supplementary Figures for a reproduction of Figure 1 using other aggregation summary methods and classification algorithms.

**Figure 1:**
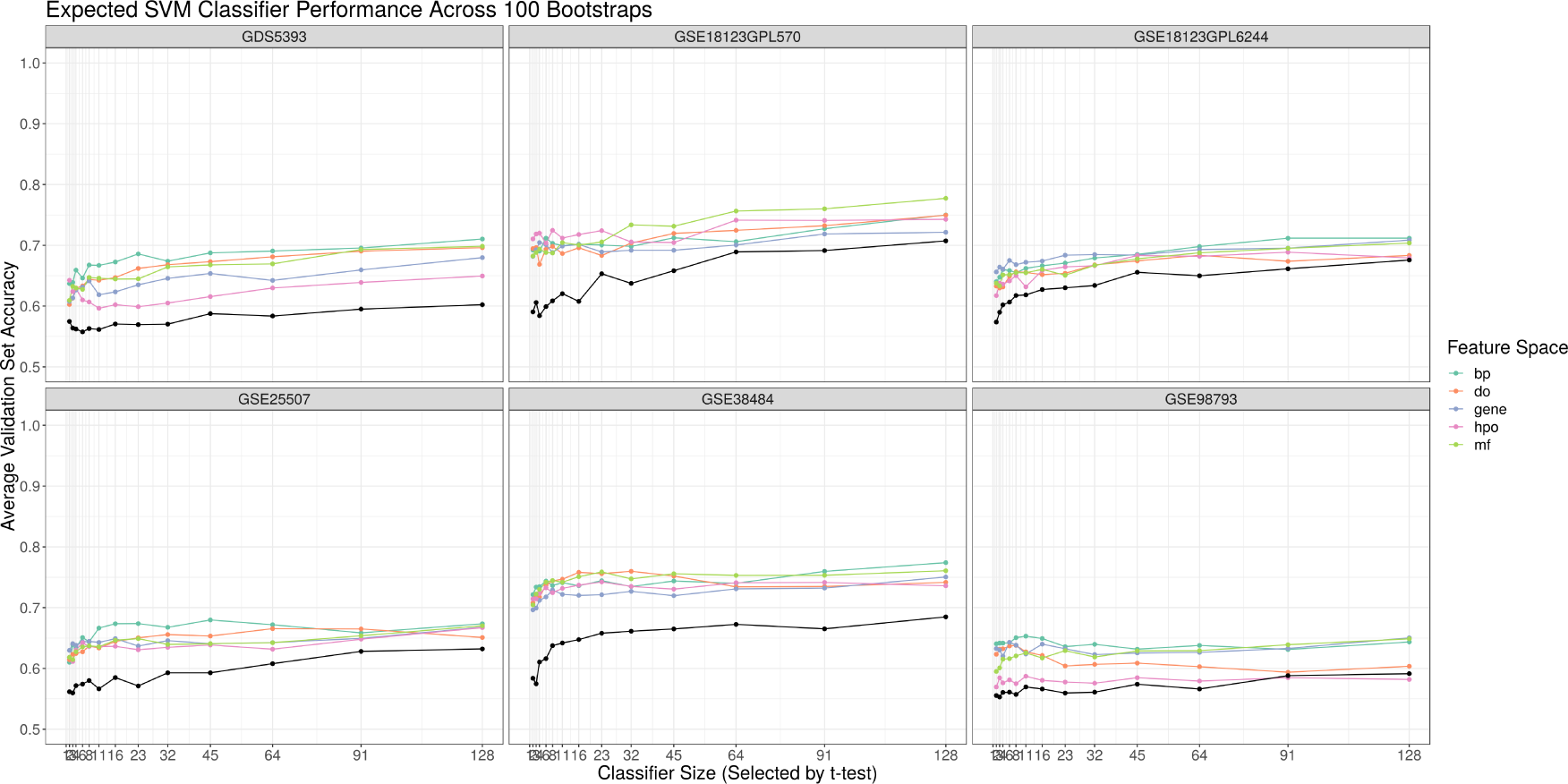
This figure shows the average Monte Carlo cross-validation accuracies for SVM classifiers (y-axis) built with the top *N* features (x-axis) across five feature spaces (gene-level or annotation-level) and six data sets (facet). Classifiers were built using features selected by a *t*-test. The black lines show baseline classifier performances for a set of randomly selected genes.

**Figure 2:**
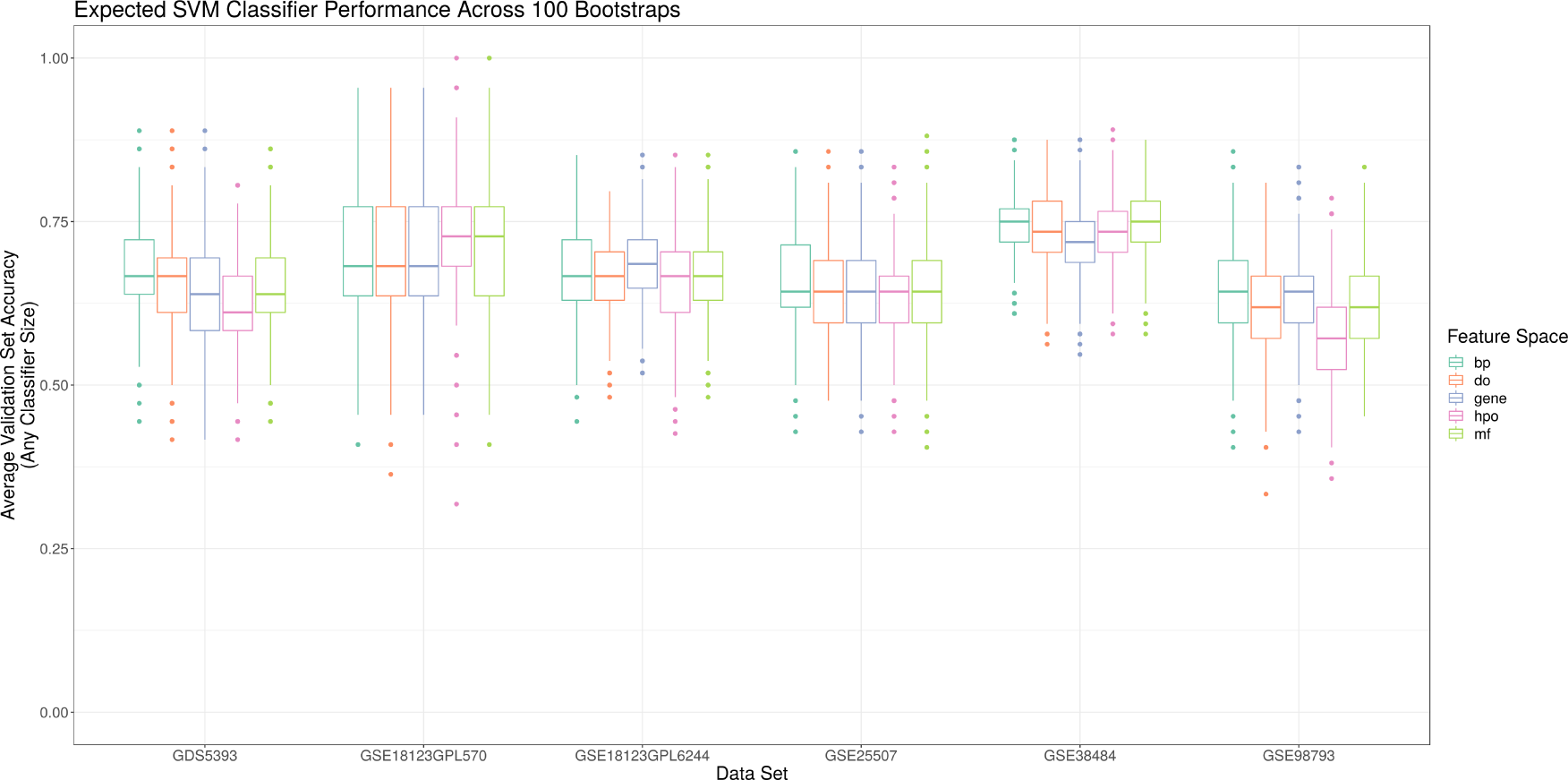
This figure shows the average Monte Carlo cross-validation accuracies for SVM classifiers (y-axis) across six data sets (x-axis) built with the top *N* features of five feature spaces (gene-level or annotation-level). Classifiers were built using features selected by a *t*-test. Boxplots pool results irrespective of classifier size.

### 3.2. Annotations as features improve stability

When assembling a new classification pipeline, it is important to know not only how well the resultant classifier will perform, but also whether its design will be robust to random differences in the training set. Since our classification pipeline depends on feature selection to prioritize features for model building, we assess this “robustness” as stability by measuring the concordance of selected features across 100 bootstraps of the training set. We consider two measures of concordance: the Baroni-Urbani and Buser Overlap for the top quartile of ranked features (**BUB25**) and the Spearman’s Rank Correlation Coefficient for all ranked features (**RHO100**). For both measures, an average concordance score of 1 suggests that the features are equivalently ranked across 100 unique cuts of the data sets, while a score of 0 suggests that the individual feature rankings never actually agree.

Figure 3 shows a box plot of all **BUB25** scores for each combination of feature space and data set (of top quartile features selected by *t*-test). Figure 4 shows a box plot of all **RHO100** scores for each combination of feature space and data set (of all features selected by *t*-test). Both figures show that that annotation-based feature spaces have more stability than the gene-based feature space (*p < .*05 by *t*-test, see Tables 4 and 5). Tables 4 and 5 present the mean differences between **BUB25** and **RHO100** scores for each feature space across all data sets bootstraps, respectively. Table 6 shows that average **BUB25** and **RHO100** scores for each feature space, and their standard deviations. We refer the reader to the Supplementary Figures for a reproduction of Figures 3 and 4 using randomly sampled features which demonstrate empirically that these concordance measures have equivalent null distributions.

**Table 4:** This table shows the 95% confidence intervals for the mean differences of **BUB25** scores between mean-based and SVM-based annotation-level classifiers (built with the Gene Ontology Biological Process (**BP**) and Molecular Function (**MF**), Disease Ontology (**DO**), and Human Phenotype Ontology (**HPO**) databases), and gene-level classifiers. We found that annotation-based feature spaces are more stable than the gene-based feature space.

|  | bp vs. | do vs. | gene vs. | hpo vs. | mf vs. |
| --- | --- | --- | --- | --- | --- |
| bp | — | -0.00340 to -0.00014 | -0.022 to -0.019 | -0.00073 to 0.00291 | -0.0036 to -0.0002 |
| do | 0.00014 to 0.00340 | — | -0.020 to -0.017 | 0.0011 to 0.0046 | -0.0017 to 0.0015 |
| gene | 0.019 to 0.022 | 0.017 to 0.020 | — | 0.020 to 0.023 | 0.017 to 0.020 |
| hpo | -0.00291 to 0.00073 | -0.0046 to -0.0011 | -0.023 to -0.020 | — | -0.0048 to -0.0012 |
| mf | 0.0002 to 0.0036 | -0.0015 to 0.0017 | -0.020 to -0.017 | 0.0012 to 0.0048 | — |

**Table 5:** This table shows the 95% confidence intervals for the mean differences of **RHO100** scores between mean-based and SVM-based annotation-level classifiers (built with the Gene Ontology Biological Process (**BP**) and Molecular Function (**MF**), Disease Ontology (**DO**), and Human Phenotype Ontology (**HPO**) databases), and gene-level classifiers. We found that annotation-based feature spaces are more stable than the gene-based feature space.

|  | bp vs. | do vs. | gene vs. | hpo vs. | mf vs. |
| --- | --- | --- | --- | --- | --- |
| bp | — | 0.0063 to 0.0134 | -0.080 to -0.073 | -0.0021 to 0.0056 | -0.0091 to -0.0017 |
| do | -0.0134 to -0.0063 | — | -0.089 to -0.083 | -0.0118 to -0.0045 | -0.019 to -0.012 |
| gene | 0.073 to 0.080 | 0.083 to 0.089 | — | 0.074 to 0.081 | 0.067 to 0.074 |
| hpo | -0.0056 to 0.0021 | 0.0045 to 0.0118 | -0.081 to -0.074 | — | -0.0109 to -0.0034 |
| mf | 0.0017 to 0.0091 | 0.012 to 0.019 | -0.074 to -0.067 | 0.0034 to 0.0109 | — |

**Table 6:**
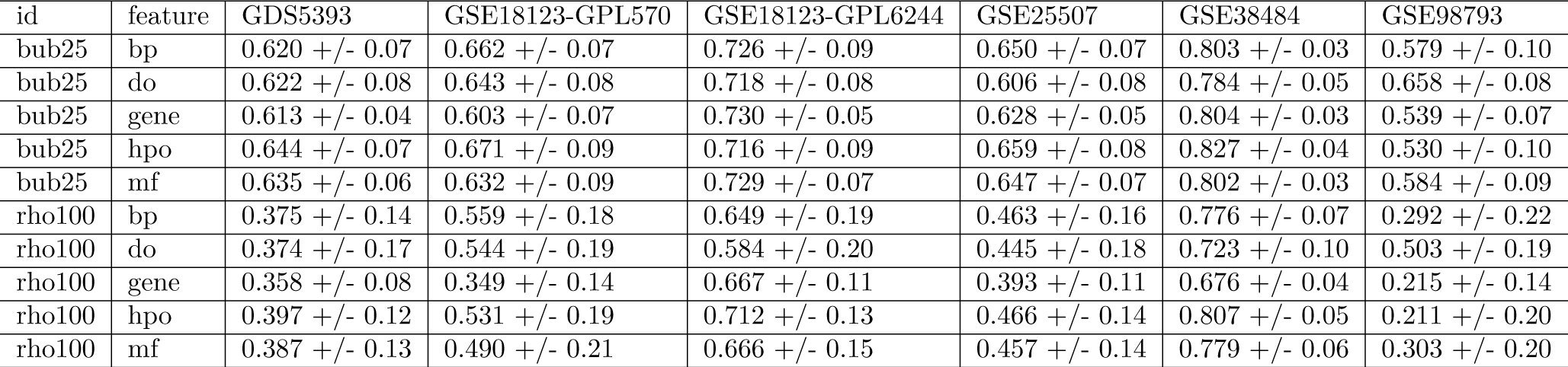
This table shows that average **BUB25** and **RHO100** scores for each feature space, and their standard deviations.

| id | feature | GDS5393 | GSE18123-GPL570 | GSE18123-GPL6244 | GSE25507 | GSE38484 | GSE98793 |
| --- | --- | --- | --- | --- | --- | --- | --- |
| bub25 | bp | 0.620 +/- 0.07 | 0.662 +/- 0.07 | 0.726 +/- 0.09 | 0.650 +/- 0.07 | 0.803 +/- 0.03 | 0.579 +/- 0.10 |
| bub25 | do | 0.622 +/- 0.08 | 0.643 +/- 0.08 | 0.718 +/- 0.08 | 0.606 +/- 0.08 | 0.784 +/- 0.05 | 0.658 +/- 0.08 |
| bub25 | gene | 0.613 +/- 0.04 | 0.603 +/- 0.07 | 0.730 +/- 0.05 | 0.628 +/- 0.05 | 0.804 +/- 0.03 | 0.539 +/- 0.07 |
| bub25 | hpo | 0.644 +/- 0.07 | 0.671 +/- 0.09 | 0.716 +/- 0.09 | 0.659 +/- 0.08 | 0.827 +/- 0.04 | 0.530 +/- 0.10 |
| bub25 | mf | 0.635 +/- 0.06 | 0.632 +/- 0.09 | 0.729 +/- 0.07 | 0.647 +/- 0.07 | 0.802 +/- 0.03 | 0.584 +/- 0.09 |
| rho100 | bp | 0.375 +/- 0.14 | 0.559 +/- 0.18 | 0.649 +/- 0.19 | 0.463 +/- 0.16 | 0.776 +/- 0.07 | 0.292 +/- 0.22 |
| rho100 | do | 0.374 +/- 0.17 | 0.544 +/- 0.19 | 0.584 +/- 0.20 | 0.445 +/- 0.18 | 0.723 +/- 0.10 | 0.503 +/- 0.19 |
| rho100 | gene | 0.358 +/- 0.08 | 0.349 +/- 0.14 | 0.667 +/- 0.11 | 0.393 +/- 0.11 | 0.676 +/- 0.04 | 0.215 +/- 0.14 |
| rho100 | hpo | 0.397 +/- 0.12 | 0.531 +/- 0.19 | 0.712 +/- 0.13 | 0.466 +/- 0.14 | 0.807 +/- 0.05 | 0.211 +/- 0.20 |
| rho100 | mf | 0.387 +/- 0.13 | 0.490 +/- 0.21 | 0.666 +/- 0.15 | 0.457 +/- 0.14 | 0.779 +/- 0.06 | 0.303 +/- 0.20 |

**Figure 3:**
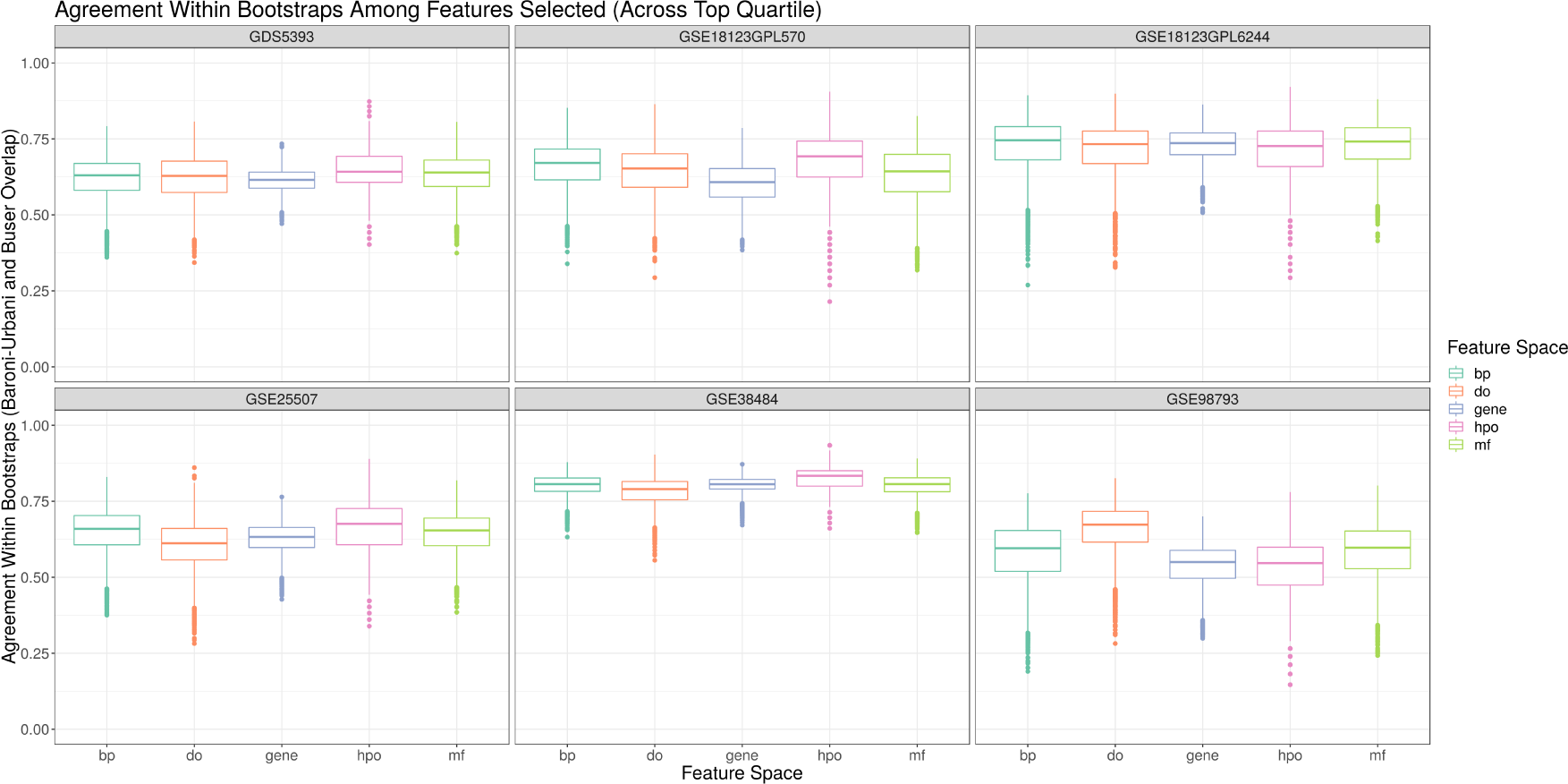
This figure shows the **BUB25** scores (y-axis) of *t*-test selected features from each feature space (x-axis) and data set (facet). For a random sample of features, the **BUB25** score has equal means irrespective of feature space.

**Figure 4:**
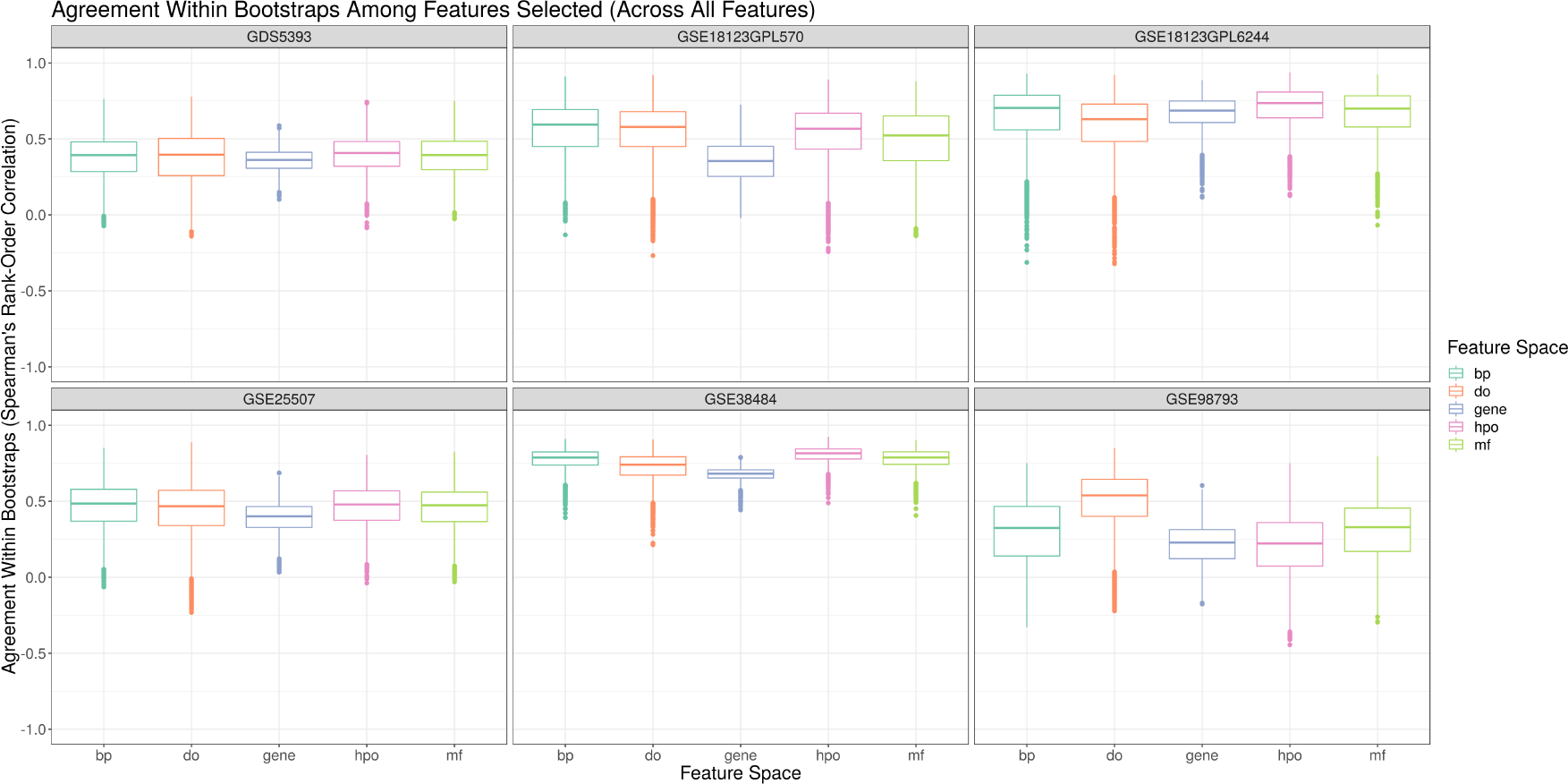
This figure shows the **RHO100** scores (y-axis) of *t*-test selected features from each feature space (x-axis) and data set (facet). For a random sample of features, the **RHO100** score has equal means irrespective of feature space.

### 3.3 Annotations as features does not improve generalizability

By using three independently collected ASD data sets (two of which comprise a single study), we can directly measure the generalized accuracy of a classifier built on one ASD data set and deployed on another. Figure 5 shows the performance of an SVM classifier trained on each data set and deployed on another (with test set positioned along the y-axis facet). Visually, we see that no approach to feature selection, regardless of the feature space, performs considerably better than randomly sampling genes. In fact, generalization is incredibly poor, especially across studies, whereby some classifiers perform no better than chance. Although annotation-based features improve cross-validation accuracies and feature selection stability, they do not improve the poor generalizability of ASD classifiers.

**Figure 5:**
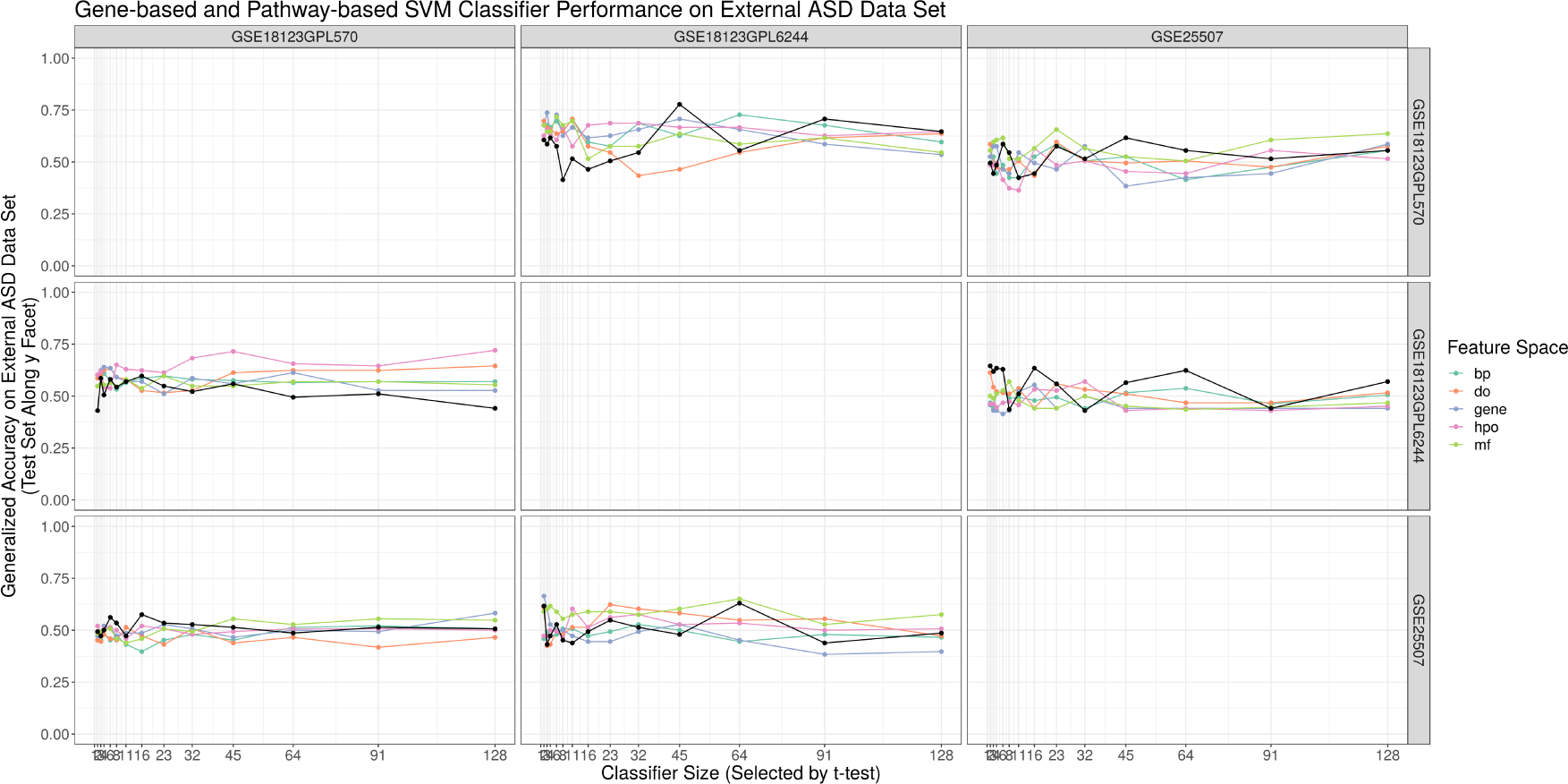
This figure shows the accuracies for SVM classifiers (y-axis) built with the top *N* features (x-axis) across five feature spaces (gene-level or annotation-level) using one ASD data set as the training set (x-axis facet) and another ASD data set as the test set (y-axis facet). Classifiers were built using features selected by a *t*-test. The black lines show baseline classifier performances for a set of randomly selected genes.

### 3.4 Annotations as features capture gene information

If we intend to interpret annotation-based classifiers directly, we posit that the annotation feature space should meaningfully represent the gene feature space. To this end, we investigated whether the selected annotation features capture the same information as the selected gene features by measuring the overlap of each “gene member set” with its corresponding gene set. Figure 6 shows the “information capture” for the four annotation-level feature spaces across the three ASD data sets. Here, we see that the **BP** feature space performs consistently well, as evidenced by the small *p*-values that suggest good agreement between the annotation and gene sets. Indeed, the principle of “information capture” seems important not only for classifier interpretation, but also classifier performance. Figure 7 shows that classifiers built using annotation feature sets with better “gene member set” overlap perform significantly better (*p < .*05 by *t*-test).

**Figure 6:**
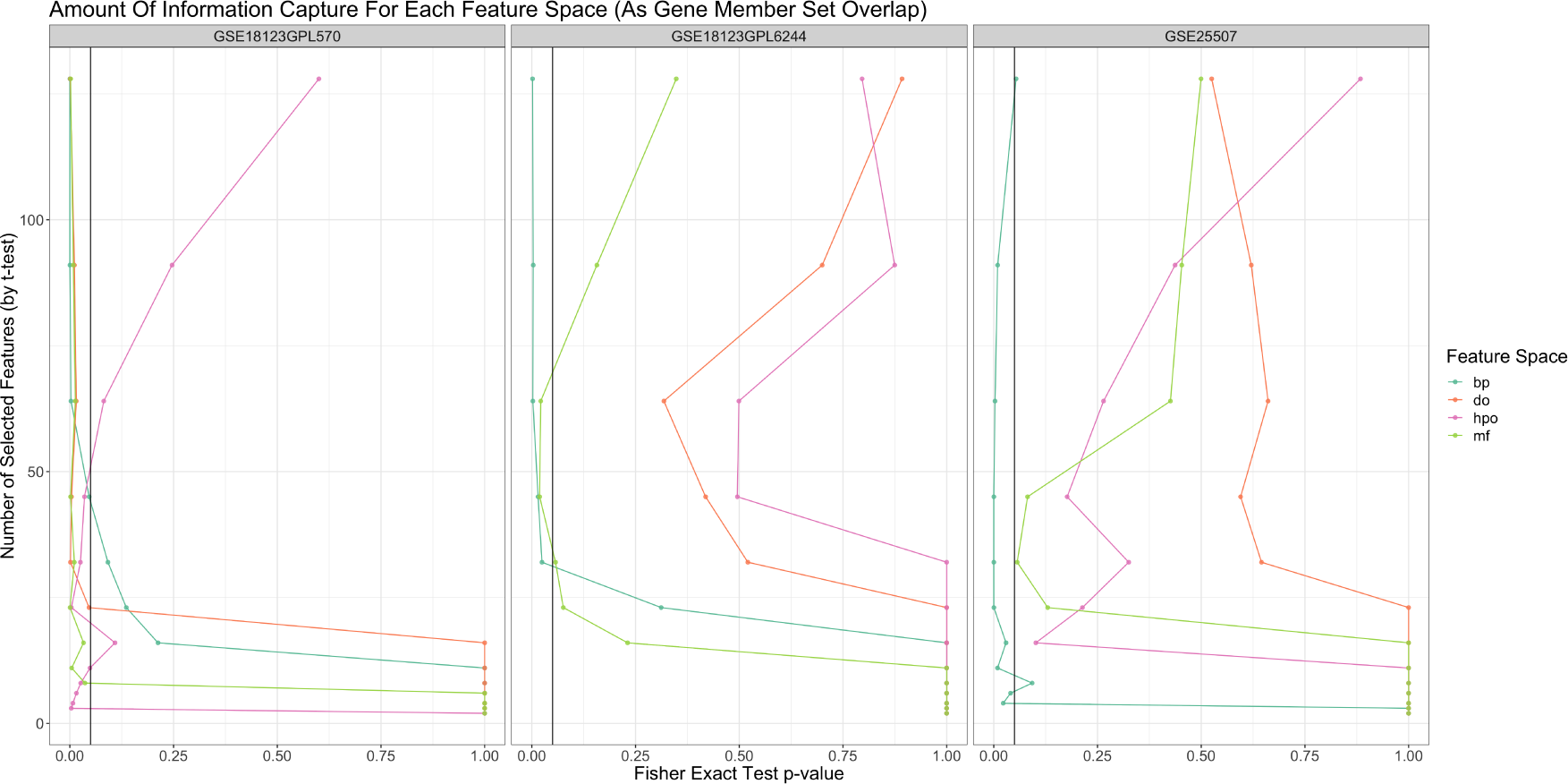
This figure shows the *p*-value of “gene member set” and get set overlap (x-axis) for the top *N t*-test selected features (y-axis) across four annotation-level feature spaces and three ASD data sets (facet). We found that annotation-based feature spaces, especially the Gene Ontology Biological Process annotation feature space, captures similar biological information to their corresponding gene-based feature spaces. The black lines indicate a *p*-value of *α* = .05.

**Figure 7:**
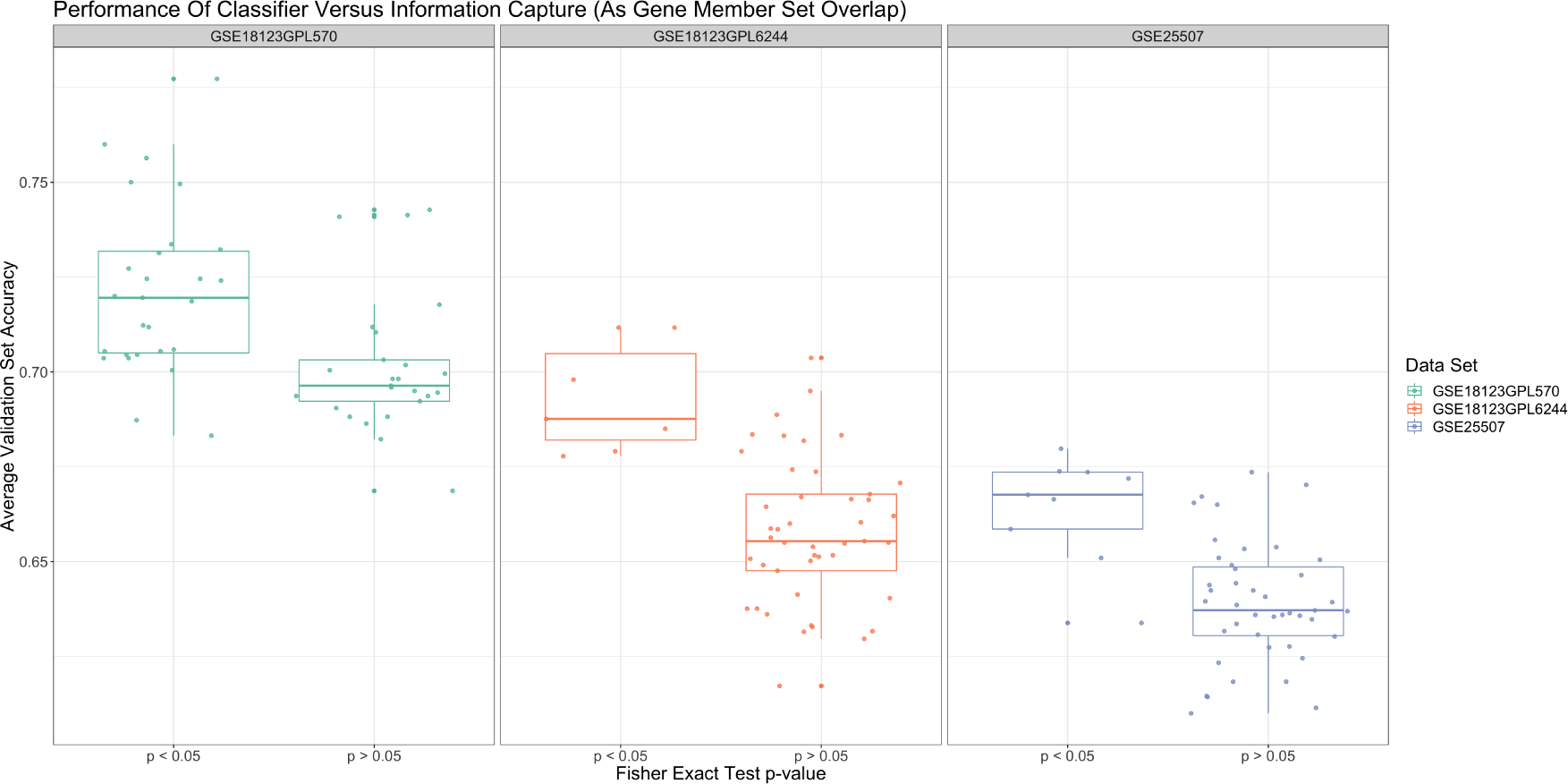
This figure shows the Monte Carlo cross-validation accuracy for SVM classifiers (y-axis) built with an annotation-based feature space having significant “information capture” or not (x-axis) across four annotation-level feature spaces and three ASD data sets (facet). For each ASD data set, accuracies are higher for those classifiers that have significant “information capture” (*p < .*05 by *t*-test). Boxplots pool results irrespective of annotation-based feature space.

## 4 Discussion

The classification of biological outcomes based on gene expression data is a growing area of research (reviewed thoroughly by [17] and [28]), where the ability to make accurate predictions from blood transcriptomes could serve as a non-invasive and clinically useful diagnostic test. Such tests would prove especially valuable to the field of neuropsychiatry, where diagnostic criteria do not have a strong molecular foundation, and where early diagnosis can improve patient outcomes (as previously demonstrated for children with autism spectrum disorders (ASD) [9]). Machine learning techniques, especially artificial neural nets (ANN) and support vector machines (SVM) [12], have grown in popularity, and have been used successfully for classifying neuropsychiatric conditions based on neuro-imaging [36] and gene expression [35].

Typically, blood-based classifiers are built using gene expression as measured by microarray or next-generation sequencing. Genes included in a classifier are described as biomarkers, and investigators often wish to know the functional role of these biomarkers (e.g., based on gene set enrichment testing of annotations from established ontological databases [31]). For this study, we hypothesized that using annotations as the features themselves would improve the classification of neuropsychiatric conditions, while providing a clearer interpretation to the researcher. Indeed, we found that annotation-based classifiers were more accurate and more stable than gene-based classifiers, although these gains were marginal. Specifically, we found that aggregating gene features into annotation features based on the mean (or sum) of the many-to-many mappings significantly improved Monte Carlo cross-validation accuracy across six data sets. Of the five feature spaces tested, we found that the Gene Ontology Biological Process (**BP**) annotation feature space outperformed others in terms of accuracy, stability, and information capture. We also found that SVM outperformed other methods (consistent with others [25]).

Although we observed some modest improvements in classifier accuracy and stability, we had expected overall larger gains. We offer a few suggestions as to why our annotation-based classifiers did not greatly outperform the gene-based classifiers. First, annotation databases are based on available experimental evidence and thus incomplete. As such, mapping gene-level expression to annotation-level expression always results in a loss of information. Second, annotation databases can draw from studies on non-human subjects, specific tissues, or cell lines. Such evidence may not meaningfully organize the gene expression of clinical blood samples. Third, this study only considered simple aggregations of gene-level expression (i.e., based on summary statistics). We did not test complex feature engineering methods as an alternative (e.g., [8, 37]). Fourth, this study only considered univariate feature selection methods. By design, these methods may select redundant annotations which individually predict the outcome accurately, but jointly add no value to the classifier. In light of these limitations, we believe that the modest improvements reported here justify further exploration into the use annotation-based features for the classification of transcriptomic data.

Although we did find that annotation-based classifiers were accurate and stable, we did not see the improvement in generalizability that we expected. Instead, we observed low generalizability for all classifiers in general. We suggest a few reasons for why our ASD models did not generalize across studies. First, the ASD label encompasses a broad spectrum of phenotypes (possibly representing a broad spectrum of aetiology). If each study recruited patients with different phenotypes, the classifier could overfit phenotype-specific signatures. Second, study differences with regard to the prevalence of medical co-morbidities could confound the ASD label. Third, study differences with regard to technical processes, including the use of different microarray platforms and data pre-processing methods, could place patient samples on an incongruous scale. Fourth, the Kong et al. data measured whole blood expression while the Alter et al. data measured lymphocyte expression, and cell type composition could confound the ASD signature [38]. Finally, it is possible that some of the intra-study accuracy observed in ASD classification is driven by the presence of confounding batch effects not present in other studies. Whatever the cause, it is apparent that ASD biomarker signatures do not reproduce across studies, as further evidenced by inconsistencies in the differential expression (DE) analysis of ASD transcriptomes. Indeed, one meta-analysis of ASD data sets found no overlap among significant DE genes across all studies [6], while another found that, even among genes significantly DE by meta-analysis, none showed even nominal DE across all studies [19]. The inability to identify consistent ASD biomarkers remains a major barrier to translating machine learning methods into clinical practice, but the problem is not apparently resolved by using annotation-based classifiers.

Although the primary purpose for building classifiers is to predict outcomes accurately, classifiers also enable an understanding of the data through the post-hoc evaluation of trained models. We believe that the use of annotation-based features better facilitates classifier understanding because it represents the data in a space that captures how scientists conceptualize biology. The annotations used here, notably **BP** and **MF**, form the foundation of gene set enrichment analysis, the most popular method for determining the biological importance of disorder-associated biomarkers. The principles of gene set enrichment analysis forms the foundation of virtually all downstream transcriptomic analyses (including DE analysis [31] and gene co-expression analysis [16]). Indeed, it is also used to assess the biological importance of features within classifiers, constituting part of the study from which the Kong et al. data derive [15]. By using annotation-based classifiers, the biological importance of the features are the features themselves.

## 5 Conclusion

In summary, we found that using annotations to engineer features improves classification accuracy and stability across six neuropsychiatric blood-based data sets. Through systematically bench-marking a bias-free classification pipeline, we found that the Gene Ontology Biological Process (**BP**) annotation feature space improves classifier performance in terms of accuracy and stability. We also noted that the top ranked annotations tend contain the top ranked genes, suggesting that the most predictive annotations are a superset of the most predictive genes. Based on this, and the fact that these annotations are otherwise used routinely to assign biological importance to genetic data, we recommend transforming gene-level expression into annotation-level expression prior to classification. We hypothesize that further research into annotation-based classifiers, especially with regard to multivariate or embedded feature selection, could result in even greater improvements to the blood-based classification of neuropsychiatric conditions.

